# HBV rewires liver cancer signaling by altering PP2A complexes

**DOI:** 10.1101/2023.03.15.532845

**Authors:** Adriana Pitea, Rigney E Turnham, Manon Eckhardt, Gwendolyn M Jang, Zhong Xu, Huat C Lim, Alex Choi, John Von Dollen, Rebecca S. Levin, James T Webber, Elizabeth McCarthy, Junjie Hu, Xiaolei Li, Li Che, Gary Chan, R. Katie Kelley, Danielle Swaney, Wei Zhang, Sourav Bandyopadhyay, Fabian J Theis, Xin Chen, Kevan Shokat, Trey Ideker, Nevan J Krogan, John D Gordan

## Abstract

Infection by hepatitis B virus (HBV) increases risk for liver cancer by inducing inflammation, cellular stress and cell death. To elucidate the molecular pathways by which HBV promotes cancer development and progression, we used affinity purification mass spectrometry to comprehensively map a network of 145 physical interactions between HBV and human host proteins in hepatocellular carcinoma (HCC). We find that viral proteins target host factors that are preferentially mutated in non-HBV-associated HCC, implicating cancer pathways whose interaction with HBV plays a role in HCC. Focusing on proteins that directly interact with the HBV oncoprotein X (HBx), we show that HBx remodels the PP2A phosphatase complex, altering its effect on tumor signaling. HBx excludes striatin-family regulatory subunits from PP2A, causing Hippo kinase activation and unmasking a requirement for mTOR complex 2 to maintain expression of the YAP oncoprotein in HCC. Thus, HBV rewires HCC to expose potentially targetable signaling dependencies.

**Significance:** Precision medicine has revolutionized cancer treatment but remains elusive for HCC. We used proteomics to define HBV/host interactions and integrated them with HCC mutations. The results implicate modifiers of HCC behavior via remodeling of host complexes and illuminate new biological mechanisms in advanced disease for therapeutic investigation.

## Introduction

Hepatocellular carcinoma (HCC) is the third leading cause of cancer death worldwide (1) and is increasing in incidence in the United States (2). Comprehensive genomic profiling of HCC has revealed only a low incidence of targetable driver mutations (3,4). Accordingly, HCC treatment is currently directed at common disease features such as angiogenesis or immune evasion (5).

The majority of HCC arises in the setting of co-morbid hepatitis due to viral infection caused by hepatitis B or hepatitis C viruses (HBV or HCV), or metabolic disorders such as non-alcoholic steatohepatitis (3,4). Some tumor-associated viruses exert direct effects on tumor maintenance, such as the degradation of *TP53* by Human Papilloma Virus (HPV) E6 (6). Hepatitis viruses primarily promote tumor initiation via increased local inflammation and cell turnover (3,4,7). The timeline for HBV- and HCV-induced tumorigenesis in murine models is consistent with an indirect role, with viral proteins causing HCC with >1 year latency (8). In contrast, HPV protein expression causes papillomatosis in mice within 2-3 months (9).

Despite its primarily indirect effect on oncogenesis, we and others have identified an HBV-associated genomic phenotype in HCC, which is observed when HBV is integrated into the host genome (3,4,10). HBV is a small, enveloped DNA virus which expresses a surface antigen (HBs), core protein (HBc), polymerase (Pol) and a putative oncogenic effector protein, HBV protein X (HBx). Pol is the only HBV protein with enzymatic activity, functioning to initiate minus strand DNA synthesis and reverse transcribe viral RNA intermediates back into DNA. HBc forms the viral capsid around the HBV genome, with surface antigen mediating hepatocyte binding (7,11). HBx has been ascribed many potential oncogenic functions, including promoting apoptosis and activating the kinase AKT (7). HBx is also necessary for HBV replication based on its ability to hijack the CRL4 E3 ubiquitin ligase (12). Earlier, members of this team reported a complete HCV protein-protein interaction (PPI) map (13). In contrast to HBx, there is no suspected HCV oncogenic effector, and few PPIs are seen with cancer-relevant proteins.

The distinct phenotype of HBV-associated HCC and putative oncogenic effects of HBV proteins motivate further study to characterize HBV’s contribution to HCC initiation and maintenance. Multilevel data integrative models are well-suited to study the complex interactions between cancer and infectious diseases. We have previously used a similar approach where we integrated host/viral PPIs with cervical and oropharyngeal cancer genomics This workflow enabled identification of cancer pathways that are modulated either by HPV in virus-associated tumors or by somatic mutations in non-viral tumors (8). Here, we extend this strategy to HBV and its impact on HCC, further advancing the analytical workflow to address the marked mutational heterogeneity of HCC. This approach yields new insights into HCC biology and demonstrates potent effects of HBV HCC genomics and tumor cell biology.

## Results

### HBV-HCC interactome

We first performed affinity purification - mass spectrometry (AP-MS) to identify HBV PPIs (6). We used a 2X Strep epitope to tag and pull down each HBV protein, including Pol, HBc and HBx. In addition, we separately examined the secreted HBc variant e antigen (HBe) and short (SHB), medium (MHB) and long (LHB) isoforms of HBs (Fig. 1B). These were overexpressed individually in HUH7 HCC cells, with AP performed using streptavidin beads. Co-purifying human proteins were identified by MS and scored with MS interaction Statistics (MiST) software (14). Based on prior publications with subsequently validated viral interactomes, we defined a MiST score threshold of 0.75, which identified 145 total high-confidence PPIs (Fig. S1A; Table S1).

**Figure 1:**
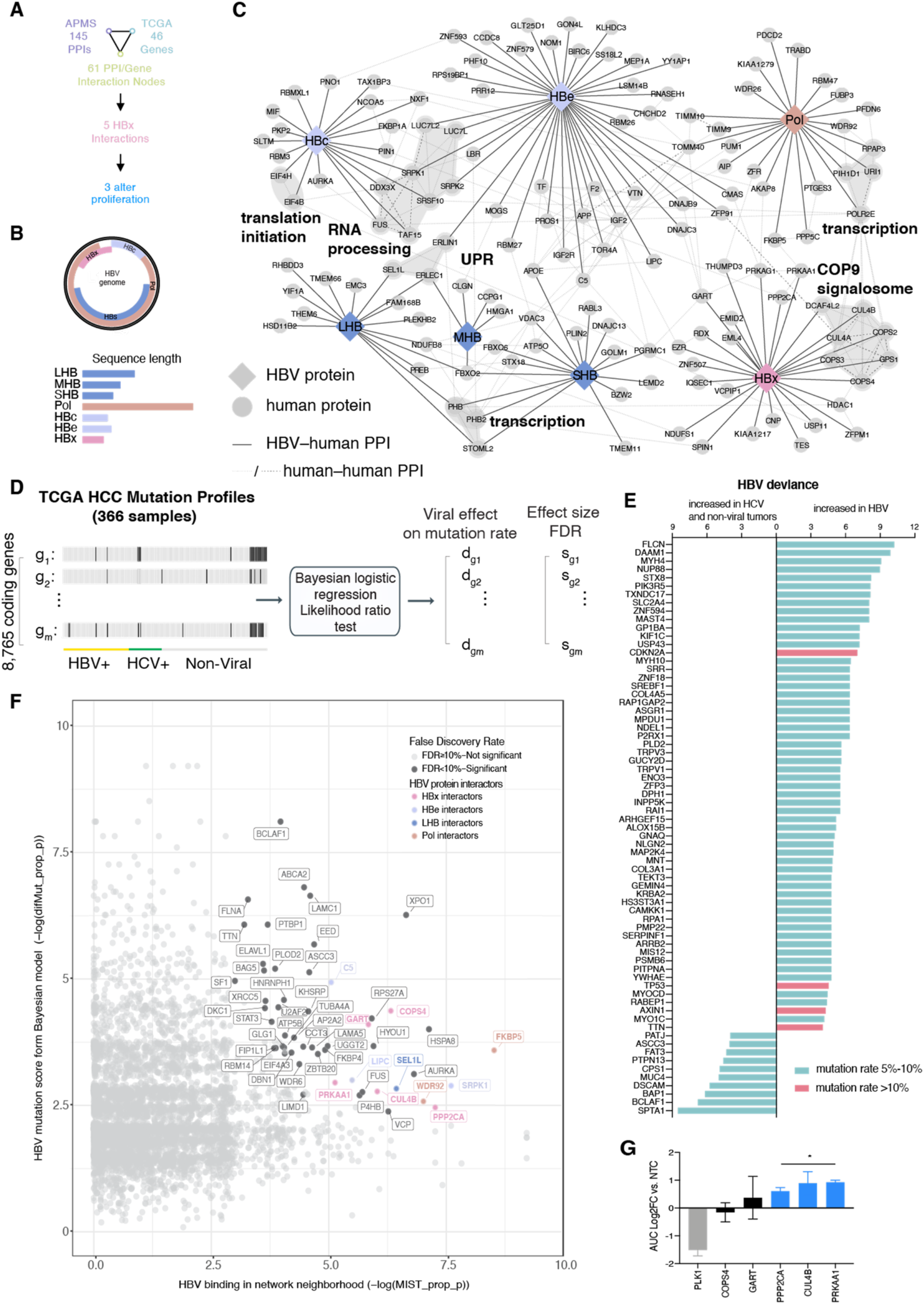
HBV-HCC protein and genomic interactomes. **A)** Overall analytical workflow: following network integration of genetic and PPI data, 5 HBx interactions were selected for further analysis with 3 altering *in vitro* proliferation. **B)** Map of the HBV genome with selected promoter sites marked, and the outline of individual genes; genes are overlapping with distinct reading frames. Individual genes are shown in linear format to show length differences of each gene tested. **C)** HBV interactome. Solid edges connect host proteins to the interacting HBV bait, while dashed lines show known human:human PPIs. Functional subsets and known protein complexes are investigator identified and designated with a grey background; n=2 biological replicates with 2 technical replicates per sample. **D)** Workflow: a Bayesian logistic regression analysis was applied to 8,765 individual non-synonymous somatic mutations identified in the TCGA LIHC project to determine the predictive effect of HBV or HCV, or either infection, on the rate of each individual mutation. Viral status was determined based on clinical annotation. **E)** Deviance in HBV and mutation rate increased in HBV (right) or increased in non-HBV (left). All genes with a mutation rate above 5% in HBV-associated or non-HBV-associated HCC, with rates between 5-10% in blue and greater than 10% shown in red. **F)** Scatter plot of propagation results: all results of network propagation are distributed by the propagated p value for relative mutation vs. HBV PPI significance. All nodes with FDR < 10% are identified, and those that interact directly with an HBV protein are color coded. **G)** Analysis of HBx PPIs that are also significant from network propagation for effects on cellular proliferation: HUH7 cells were engineered to express dCAS9-KRAB and then stably transduced with 4-5 sgRNA against each HBV PPI that reached significance in network analysis; PLK1 is included as an essential gene control. Effects on proliferation with real time microscopy over 120 hours are shown (* p < 0.05); n=3 biological replicates per sgRNA with a minimum of 3 technical replicates per sample. Area under the curve (AUC) analysis of cell count at each time point was developed and compared to cells transduced with a non-targeting control (NTC) sgRNA.

Functional analysis of the identified proteins with the Reactome database (15) revealed enrichment of proteins involved in pre-mRNA processing (HBc), nucleotide excision repair and mTOR signaling (HBx) and the unfolded protein response (HBe and LHB; Fig. S1B). The full PPI map (Fig. 1C) included novel PPIs of HBV subunits with biologically significant host proteins including eIF4H and DDX3X (HBc) as well as Prohibitin 1 and 2 (LHB, SHB). It also included many known and suspected HBx interactions such as PRKAA1, PPP2CA, HDAC1, and the CRL4 E3 ubiquitin ligase complex. Selected PPIs were confirmed by AP-western (Fig. S1C). This interaction network provides an expanded view of the potential effects of HBV in HCC cells.

### HBV significantly impacts gene mutation status in HCC

We next identified genes with differential mutational status relative to viral infection. We used Bayesian logistic regressionto identify protein-coding genes with recurrent genetic alterations in The Cancer Genome Atlas (TCGA) HCC cohort (3) for which the alteration rate was significantly dependent on the presence of HBV or HCV infection (Fig. 1D). This strategy allowed us to evaluate the impact of the two hepatitis viruses on a broad spectrum of commonly and rarely mutated genes in HCC. We identified 70 genes with differential mutation based on viral status (*p* value <0.05, FDR < 20%; Table S2). The majority of differentially mutated genes had increased mutation frequency upon HBV viral infection (Fig. 1E). No statistically significant differences were observed in HCV-associated HCC compared to the remaining subsets. Notably, the tumor suppressors *TP53* and *MAP2K4* and the oncogene *GNAQ* were preferentially mutated in HBV+HCC, while the tumor suppressor *BAP1* was preferentially mutated in non-HBV-associated HCC. These data confirm an impact of HBV on HCC genomes, but only provide partial insight into the ongoing role of HBV in HCC. Thus, we next pursued further investigation into direct effects of HBV.

### Network model integrates oncogenic HBV protein and genetic interactions

To identify PPIs with potential oncogenic functions, we used a network-based strategy to integrate HBV/host PPIs with differential HCC mutations, following the method some of us had established earlier for HPV (6). Point mutations and copy number aberrations that were less frequent in HBV-associated HCC (in comparison to HCV-associated and non-viral HCC) were used to identify mutations that might phenocopy HBV PPIs. This analysis followed the rationale that increased incidence of a mutation in the absence of HBV could indicate an important pathway in HCC that is directly modified by a PPI in HBV-associated tumors.

We applied statistical confidence measurements for each data set to the ReactomeFI network, a public catalog of 229,300 known pathway relationships among human proteins, including PPIs, protein–gene transcriptional regulatory interactions, and metabolic reactions. We then used the framework of network propagation (15). Deviances reflecting significance of differential mutation and MiST scores reflecting PPI confidence were propagated separately across the network to generate a *p* value for each protein. Propagated *p* values were combined, resulting in a set of 61 network nodes with FDR < 0.1, indicating Proximity in ReactomeFI to HCC differential mutations and to HBV/host PPIs (Fig. 1F, Table S3). This set includes genes that did not meet our confidence threshold to be considered HBV PPIs but through network proximity and/or differential mutation meet the threshold of inclusion (Fig. S2A, shown in dark grey). This analysis allowed prioritization of a subset of the HBV/host PPIs and added related genes to their local networks.

For HBc, interactions with SRPK1 and FUS were expanded to include KHSRP, ELAVL1 and PTBP1, giving a larger group of host factors in HBV’s known effects on splicing (16). Network propagation did not expand the set of PPIs for HBX. Instead, it allowed PPI prioritization based on cancer relevance, nominating COPS4, CUL4B, GART (a purine synthesis enzyme), PPP2CA and PRKAA1 as candidates for further study. Given HBx’s putative oncogenic role, we focused on these HBV PPIs and assessed their impact on cell proliferation with CRISPR interference, using the essential Polo-like kinase 1 (PLK1) as a positive control. Knockdown of *CUL4B, PPP2CA* and *PRKAA1* increased HUH7 proliferation, while knockdown of *GART* and *COPS4* did not, showing a good concordance between our systems analysis and a relevant cancer readout (Fig. 1G, S2B).

### HBx physically remodels host PP2A complexes

With these HBx/host PPIs prioritized based on their impact on cell proliferation, we next assessed the mechanism by which HBx can act on these targets. We focused on host protein phosphatase 2A (PP2A), given that it has been previously validated to impact HCC behavior in the context of HBV, but with conflicting effects in different reports (17–19). Given previous findings that HBx hijacks CRL4 by recruiting SMC5/6 for ubiquitylation (12), we hypothesized that HBx might alter PP2A’s substrate specificity.

Accordingly, we performed quantitative AP-MS with the PP2A catalytic subunit (PP2Ac, gene *PPP2CA*) in the presence or absence of HBx. The PP2A holoenzyme consists of PP2Ac, the scaffolding subunit PP2AA (gene *PPP2R1A*), and one of >15 regulatory subunits. These subunits direct PP2Ac’s substrate selection; several are tumor suppressors, including PPP2R4 and the PPP2R5 families (20). The PP2A interactome showed significant changes in its interaction with regulatory subunits in the presence of HBx (Fig. 2A; Table S4). HBx expression resulted in reduced interaction between PP2Ac and PP2A regulatory subunits B alpha and delta (*PPP2R2A, PPP2R2D*), PP2A regulatory subunit B’ delta (*PPP2R5D*), STRN3 and STRN4. STRN3 and STRN4 are components of the Hippo pathway regulating STRIPAK complex (21). HBx expression also resulted in reduced PP2Ac interaction with STRIPAK components STRIP1 and MOB4, as well as the DDR proteins RAD23A/B (Fig. 2B).

**Figure 2:**
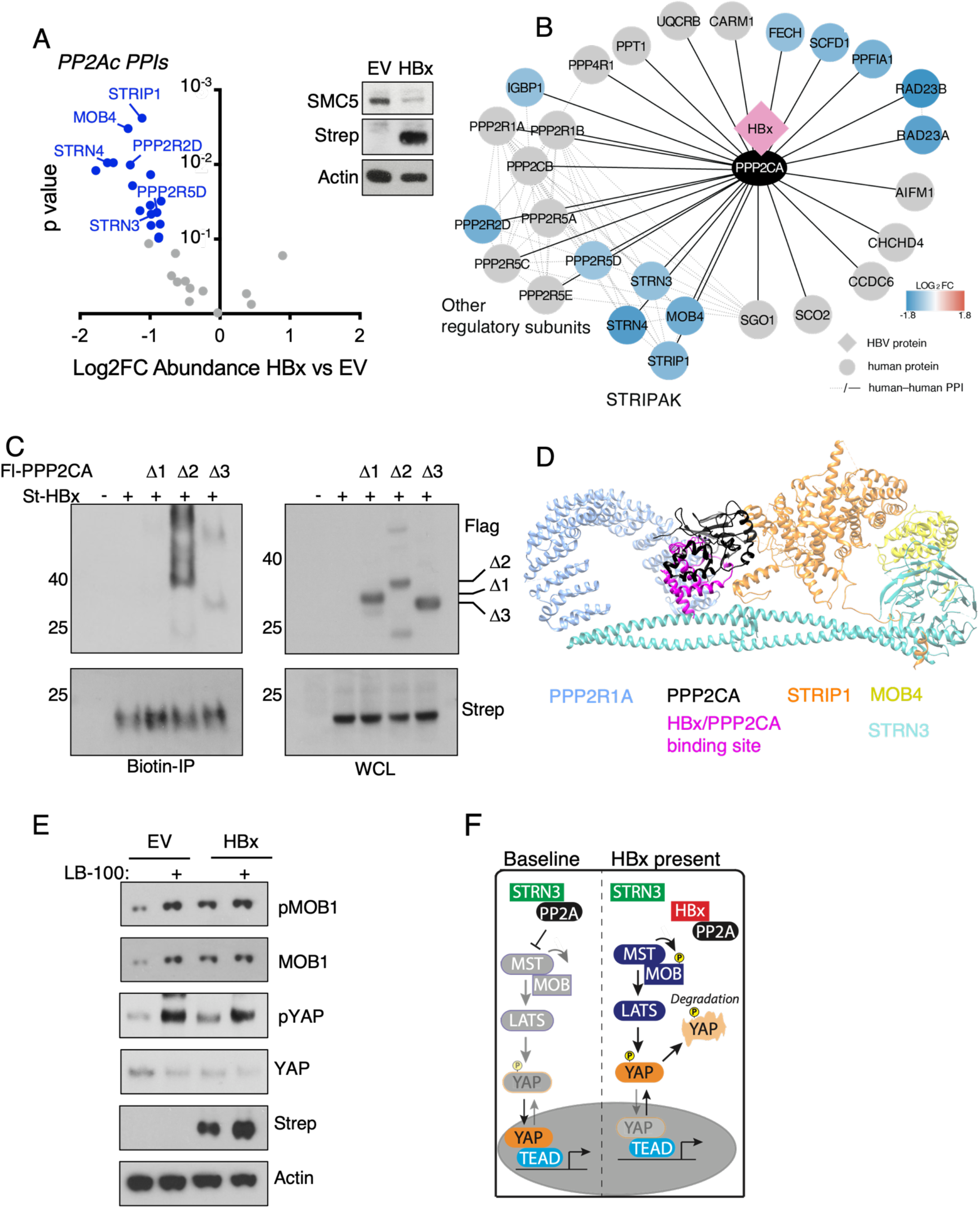
HBx remodels the PP2A holoenzyme. **A)** Physical remodeling of the PPP2CA complex in HUH7 cells with co-transfection of FL-PPP2CA and ST-HBx vs. FL-PPP2CA alone. Western blot confirmation of St-HBx expression and SMC5 knockdown along with empty vector (EV). High confidence PPIs are shown, with statistically significant changes in abundance in color; proteins of interest directly labeled. n=2 biological replicates with 2 technical replicates per sample. **B)** Cytoscape presentation of AP-MS results from A, with proteins with significantly altered abundance in color. There were no PPIs with increased abundance. **C)** Deletion mapping of the HBx/PPP2CA interaction with co-overexpression; for PPP2CAΔ1, 1-108aa are deleted: PPP2CAΔ2, 109-189aa are deleted; PPP2CAΔ3, 190-310aa are deleted. Biotin IP was used for complex purification and compared to whole cell lysate (WCL); n=2 biological replicates with 1 technical replicate per condition. **D)** Model of the HBx/PPP2CA interaction and its physical proximity to the PPP2CA/STRN3 interface overlaid onto the 7K36 cryo-EM structure. **E)** Effect of HBx expression on PP2A/STRIPAK targets: HUH7 parental cells were transfected with pCDNA4 (EV) or HBx-S overnight and then treated with PBS or 10μM LB-100 for 4 hours. n=2 biological replicates with 1 technical replicate per condition. **F)** Model of findings: when HBx is expressed, PP2A is displaced from STRIPAK, allowing increased activity of the Hippo kinases MST1/2 and LATS1/2, resulting in increased MOB phosphorylation and YAP degradation.

We used truncations to define the HBx/PP2Ac interaction surface and determined that HBx interacts with the first 108 amino acids of PP2Ac (Fig. 2C). This was modeled on the STRIPAK cryo-EM structure (22), showing that HBx binds to PP2Ac in a region that positions it to block binding of STRN3 and displace its associated complex components (Fig. 2D).

STRIPAK-associated PP2A can inhibit the Hippo tumor suppressor pathway via effects on the Hippo kinases MST1 and MST2 (21). Consistent with a loss of PP2A/STRIPAK activity, we found that HBx overexpression results in increased Hippo activation marked by phosphorylation and stabilization of the MST1/2 substrate MOB1 and phosphorylation-induced degradation of the terminal Hippo target YAP. Importantly, the same effect was seen with the direct PP2Ac inhibitor LB-100, with little additive impact of LB-100 and HBx overexpression (Fig. 2E, F).

### HBx modulates HCC signaling by disrupting PP2A effects

The interaction between HBx and PP2A suggests a mechanism by which HBV can regulate HCC signaling. Accordingly, we used global phosphoproteomics to assess changes in cellular signaling pathways in HUH7 cells upon HBx expression. Secondary analysis was performed with the Phosfate kinase attribution tool (23) uses annotated phosphopeptides to identify kinases with altered activity in the presence of HBx. These data showed HBx-induced changes in motility, cell cycle/DNA damage, and stress and MAPK/PI3K pathways (Fig. 3A; Table S5,6), including a previously described increase in AKT activity (7). Pro-motility signaling such as Myosin lightchain kinase (MYLK) and p21-activated kinase 1 (PAK1) were also regulated. These are also known targets of STRIPAK-associated PP2A (21). We tested a subset of targets by western blot (Fig. 3B). As seen with the Hippo kinases, HBx or LB-100 increased PAK1 autophosphorylation, albeit mildly. We also confirmed that HBx overexpression increases AKT activity, while LB-100 abolishes AKT activation in the presence or absence of HBx, suggesting that PP2A is necessary to maintain AKT activity.

**Figure 3:**
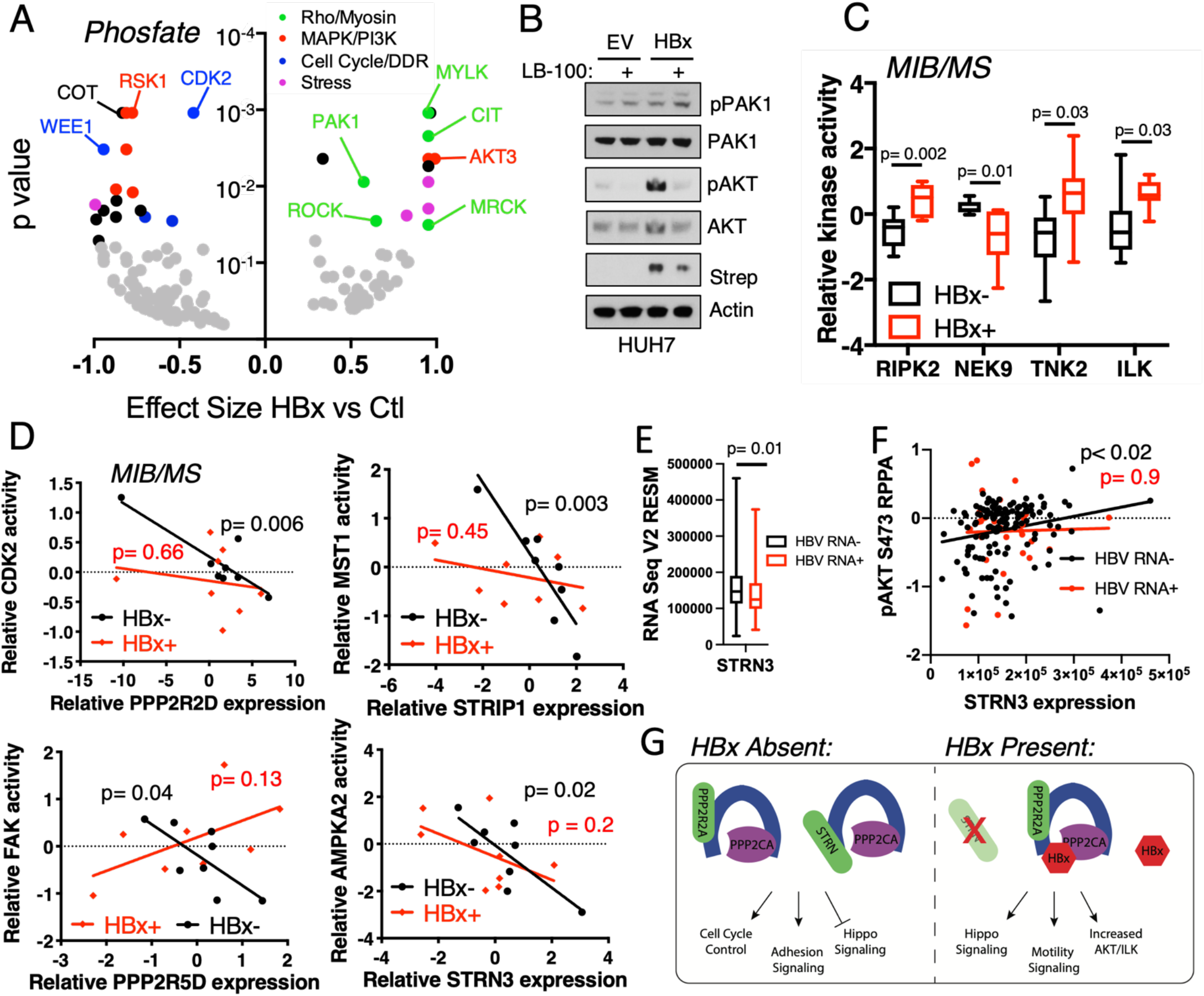
HBx alters HCC signaling by disrupting PP2A effects. **A)** Volcano plot of significant alterations in imputed kinase activation based on Phosfate analysis of global phosphoproteomics of HUH7 cells with doxycycline-induction of HBx for 48 hours. **B)** Effects of HBx overexpression or 4 hours treatment with 10μM LB-100 on imputed HBx-modified targets. **C)** Significantly altered kinase activity across a panel of 16 HCC lines, including 8 with detectable HBx mRNA expression; kinase activity determined using multiplex inhibitor beads (MIBs)/mass spectrometry (MS). p values derived using t tests. **D)** Correlation between PP2A component expression and individual kinase activity in HBx+ or HBx-cell lines. **E)** Expression of STRN3 in HBV RNA- vs. HBV RNA+ HCC specimens from TCGA where there was also RPPA data available. **F)** Correlation between STRN3 mRNA expression and AKT S473 phosphorylation integrating TCGA RNASEQ and RPPA data. **G)** Overall model for HBx effects on signaling: In the absence of HBx expression, PP2A function is determined by its typical upstream regulation mechanisms as well as the overall expression of its regulatory subunits (left). However, when HBx is present, its interaction with PPP2CA selectively disrupts PP2A function towards targets with effects on PI3K/mTOR signaling, adhesion/motility and the Hippo tumor suppressor pathway (right).

To understand signaling in the context of basal HBx expression levels, we assessed a panel of liver cancer cell lines from HBV positive (n=11) and negative (n=5) patients. HBx gene expression was assessed by qPCR, with which we found that 8 of 11 HBV-associated HCC lines had detectable HBx levels (Fig. S3A). As control, we used PathSeq (10) to identify HBV sequences in RNA-Seq data available for 11 of these lines and found perfect concordance with our qPCR results.

We then performed kinome profiling using Multiplex Inhibitor Beads coupled with mass spectrometry (MIB/MS) to compare kinase expression and activity in HCC cells (Table S7). As MS detection is influenced by kinase expression levels, MIBs have somewhat reduced detection of kinases with low abundance but favor the measurement of kinases with poorly annotated targets (24). In contrast to the isogenic system used for global phosphoproteomics analysis, the heterogeneity between cells in this larger collection resulted in relatively few consistently altered kinases based on HBx expression. Notably, these included the pseudokinase ILK, which is anticipated to still be detectable by MIBs via allosteric regulation of an ATP binding site (25), as well as the pro-inflammatory kinase RIPK2 and the pro-motility kinase TNK2 (Fig. 3C). ILK was of particular interest as it signals in concert with AKT and mTOR complex 2 (mTORC2) (26).

This dataset allowed us to assess whether endogenous HBx expression might alter PP2A activity. We studied this relationship by correlating expression levels of PP2A regulatory subunits whose presence in the holoenzyme is blocked by HBx (Fig. 2B) with signaling of their known targets. Importantly, there was also no significant difference in expression of these subunits in HBx and non-HBx expressing cell lines (Fig. S3B). As expected, we observed consistent negative correlation between regulatory subunit expression and kinase activity in non-HBx expressing cells, including PPP2R2D with CDK2, PPP2R5D with FAK, and AMPKA2 with STRN3 (Fig. 3D). STRIP1 is a STRN3 interacting protein that is present in MST1/2 regulating PP2A complexes (27). Expression of STRIP1 was significantly correlated with MST1/2 activity in non-HBx expressing cells (Fig. 3D), consistent with previous results (21). However, there was no correlation between PP2A subunit expression and activity of these kinases in HBx-expressing lines.

We next used data from the TCGA to connect our findings to *in vivo* HCC behavior. Stratifying tumors by HBV RNA expression as described previously (10), we noted that STRN3 is upregulated in non-HBV RNA expressing HCC, consistent with its role in driving mitogenic effects of PP2A (Fig. 3E). STRN3/PP2A containing complexes have been shown to increase AKT activity (28). Consistent with our *in vitro* data and previous reports, we saw that STRN3 expression correlated with increased AKT pS473 by reverse phase protein array (RPPA) in the absence of HBV RNA, whereas AKT phosphorylation was decoupled from STRN3 levels in the presence of HBV RNA (Fig. 3F). Of note, basal AKT activation was not significantly different with or without HBV RNA expression. mRNA expression of those PP2A regulatory subunits that are not influenced by HBx (e.g. PPP2R5A) still correlated with AKT phosphorylation (not shown). These data further supported our model that HBx shifts PP2A substrate specificity through its interaction with regulatory subunits, thus influencing key HCC signaling pathways including AKT and Hippo (Fig. 3G).

### HBx effects on PP2A unmask mTOR Complex 2 regulation of YAP in HCC

Although signaling can be considered as a series of linear kinase relays, most pathways exist in complex interconnected networks. Thus, the upregulation of a tumor suppressive kinase pathway suggests that another aspect of HBx-induced signaling remodeling may exert an opposing effect to maintain YAP expression. For this reason, we analyzed the impact of AKT and mTOR inhibition on YAP stability in HBx-expressing cell models.

Consistent with results in HUH7, HBx overexpression and LB-100 treatment both reduced baseline YAP levels in Hep3B. Cells were also treated with the allosteric AKT inhibitor MK2206 and the ATP-competitive mTORC1/mTORC2 mTOR inhibitor TAK-228 (sapanisertib). These data showed reduced YAP protein levels with each compound, with TAK-228 having the greatest effect. All effects were more pronounced in the presence of HBx (Fig. 4A). Also, we used CRISPRi to reduce expression of the PSK ILK, finding that it decreased YAP levels independent of HBx levels (Fig. S4A).

**Figure 4:**
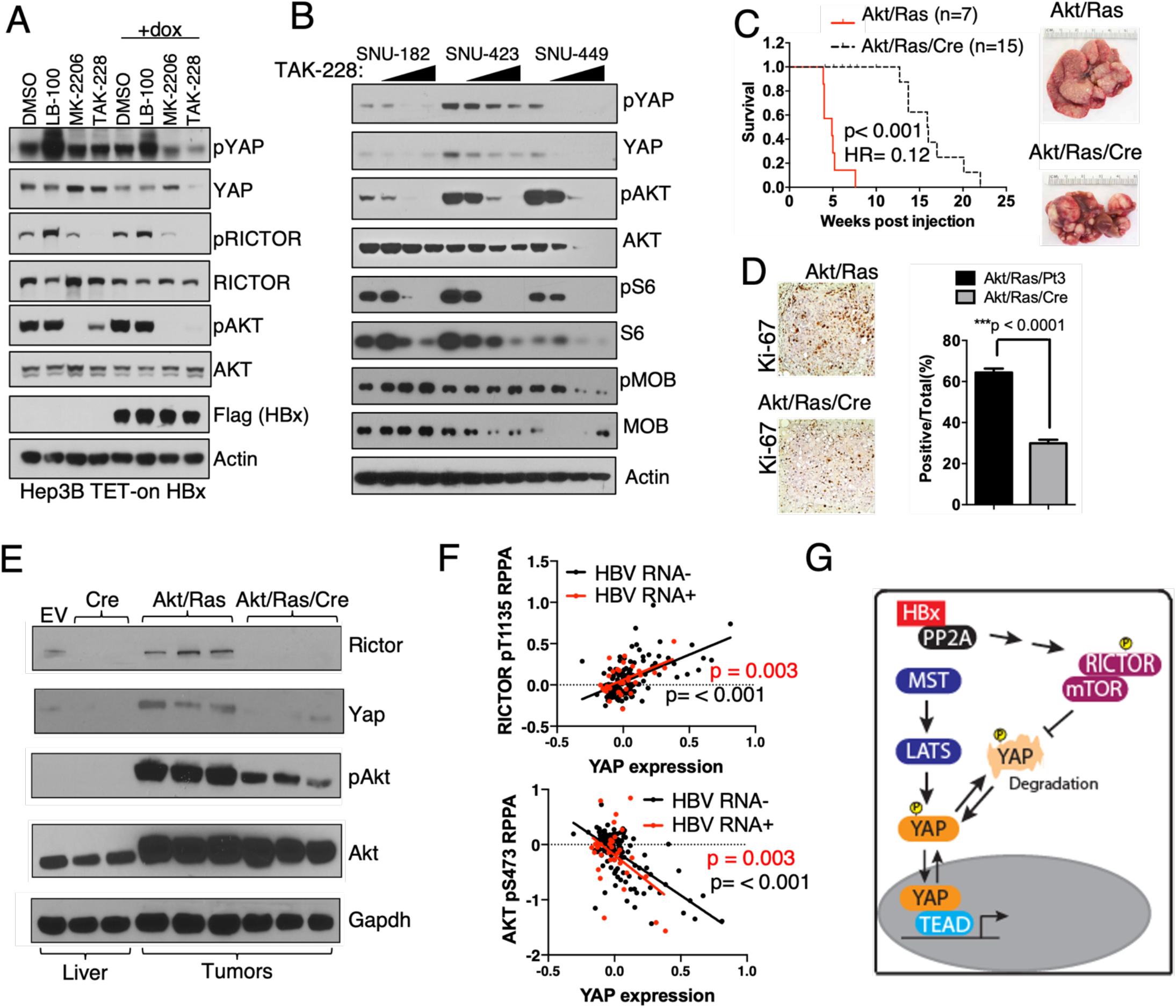
HBx stabilization of YAP via mTORC2. **A)** Impact of HBx on regulation of YAP expression by AKT and mTOR. Hep3B cells with tet-inducible FL-HBx were treated with dox for 48 hours and then 1mM MK-2206 or 100nM TAK-228 or 10μM LB-100 for 4 hours. n=2 biological replicates with 1 technical replicate per condition. **B)** Effect of DMSO or 20, 100 or 500 nM TAK-228 treatment for 24 hours on YAP expression in HBV+/HBx+ SNU-182 and SNU-449 HCC lines or HBx-SNU-423 line assessed by immunoblot. n=3 biological replicates with 1 technical replicate per condition. **C)** Kaplan-Meyer survival analysis of *Rictor^fl/fl^* mice following hydrodynamic tail vein injection with Akt, Ras and empty vector or Akt/Ras/Cre. Akt/Ras/Vector mice were sacrificed at 5 weeks because of evident morbidity, and Akt/Ras/Cre sacrificed over time up to 22 weeks. Characteristic images of livers at necroscopy. **D)** Representative images and quantified Ki-67 staining in Akt/Ras/Vector and Akt/Ras/Cre mice. **E)** Immunoblot analysis of Rictor, Yap, mTORC2 target Akt S473 phosphorylation in *Rictor^fl/fl^* livers transduced with empty vector or Cre, and then Akt/Ras or Akt/Ras/Cre tumors macrodissected at necroscopy. n=2 biological replicates with 3 technical replicate per condition. **F)** Pearson correlation between RICTOR pT1135 or AKT pS473 and total YAP protein levels in HBV-associated and non-HBV associated HCC by RPPA from the TCGA LIHC project. **G)** Summary of model, showing parallel effects of HBx to activate Hippo and mTORC2 signaling to maintain YAP expression.

These data suggested that ILK/AKT/mTORC2 mediated signaling maintains YAP expression in HCC and may be particularly important in the context of HBV infection. As AKT and ILK both signal to mTORC2, we next confirmed that TAK-228 treatment reduces YAP protein levels in the HBx-expressing cells SNU-182 and SNU-449. This effect was also seen in the non-HBx expressing line SNU 423 (Fig. 4B), which has high levels of ILK activity based on the MIBs assay (Table S7).

We confirmed that TAK-228’s effect on YAP expression was due to mTORC2 inhibition by comparing the effects of the allosteric mTORC1 inhibitor everolimus, the Rapalink mTORC1 inhibitor E-1035, and TAK-228 in SNU-182, observing that YAP levels are only reduced when mTORC2 is inhibited. Furthermore, this effect was sensitive to the proteasomal inhibitor MG-132, suggesting that mTORC2 acts on YAP degradation (Fig. S4B). Consistent with post-transcriptional YAP regulation, reduced mRNA expression of canonical YAP targets was also observed following TAK-228 treatment in cell lines, while YAP mRNA was not significantly changed (Fig. S4C).

To confirm this observation *in vivo* and validate the role of mTORC2 as a regulator of YAP in HCC, we tested the impact of *Rictor* deletion in an Akt/Ras-driven murine HCC model. Here, we found that Cre-mediated *Rictor* deletion resulted in longer survival compared to pT3 empty vector, with robust induction of aggressive tumors when Rictor was intact, improved survival and reduced proliferation in the absence of Rictor (Fig. 4C-D). Furthermore, when confirming *Rictor* deletion, we saw near elimination of YAP expression, as well as reduced but persistent Akt pS473 (Fig. 4E). These findings show that mTORC2 contributes to YAP regulation *in vivo* and again support its effect in the absence of HBx.

These data suggest that the balance between Hippo and mTORC2 signaling control YAP expression in HCC. Consistent with our hypothesis that HBx accesses important cellular pathways rather than inducing them *de novo*, mTORC2 effects on YAP but that it can be potentiated by either HBx or other factors signaling to these pathways. Consistent with this idea, we noted close correlation between levels of mTORC2 activation based on RICTOR pT1135 and YAP in HBV RNA+ and non-HBV RNA expressing specimens in TCGA. We also found that YAP and AKT pS473 were strongly inversely correlated in this data set, suggesting that our *in vitro* observation that the AKT inhibitor MK2206 reduced YAP levels was a result of AKT’s role in maintaining mTORC2 activity, rather than AKT stabilizing YAP directly (Fig. 4F). Thus, HBx induced signaling changes remodel a kinase network around the control of YAP levels, augmenting an inhibitory signal (Hippo) and creating a potentially targetable signaling vulnerability on AKT/mTORC2 (Fig. 4G).

## Discussion

This work describes the first complete HBV/HCC protein-protein and genomic interactome, and refinement of our network propagation strategy to identify cancer-relevant PPIs in genetically heterogeneous context of HCC. Our tiered analytical strategy allowed us to define remodeling of PP2A as an important HBV effector function. The increase in Hippo pathway activity resulting from HBx effects on PP2A results in a dependency on mTORC2 function to maintain YAP protein levels. The significant alterations of PP2A function and the HCC kinome in the presence of HBx suggest that this may be one of several signaling dependencies emerging in the context of HBV infection. Further, mTORC2’s maintenance of YAP expression in non-HBx-expressing specimens supports our model that HBV acts through cellular pathways that also support non-HBV-associated HCC.

This work advances the use of network strategies and multi-modality integrative models to study viral effects on cancer and reinforces the importance of protein interaction networks in defining disease-relevant genetic relationships. The mutations in non-HBV HCC that identified oncogenic HBV PPIs are not focused on a single oncogene or tumor suppressor, but rather occur across multiple members of a protein complex. By using network propagation, we were able to decode these relationships in an unbiased fashion and identify biologically meaningful relationships. This will likely broaden the utility of this strategy for the many other tumor associated viruses with indirect effects (29). Similarly, the integration of proteomic data sets to assess the impact of HBx on PP2A interaction partners and downstream signaling allowed for the identification of a clear pattern of effects in multiple cell models.

Viral disruption of PP2A function is common, with displacement of selected tumor suppressive regulatory subunits also seen in the context of the SV40 small T antigen (30). The remodeling of the PP2A complex by HBx decoupled it from the regulation of multiple cellular signaling pathways, most notably disrupting its inhibition of the Hippo tumor suppressor and decreasing YAP protein expression. This apparently growth inhibitory effect of HBx is countered by a PP2A-dependent increase in mTORC2/AKT activity, which can maintain YAP protein levels by blocking its degradation. This effect of mTORC2 occurs even in the absence of HBx and is confirmed in mouse models and RPPA data from human HCC, suggesting it is a robust mechanism with potential impact on human disease.

Although our study does not provide evidence of preferential activity of mTORC1/2 inhibitors in HBV-associated HCC models, it did suggest a connection between Hippo pathway activity and mTOR dependence. The uncovering of this new regulatory mechanism in HCC has potential implications for current therapeutics, including ongoing trials such as the Phase 2 study of the mTORC1/mTORC2 inhibitor ATG-008 in HBV-associated HCC (NCT03591965). Specific relevance of viral infection is also seen in current standard of care agents, with a marked improvement in responses to the combination of atezolizumab and bevacizumab in patients with co-morbid HBV (31). This finding may be in part due to HBV effects on AKT, known to drive angiogenesis, but could also reflect the HBc interactions with splicing proteins described above.

Finally, while our analysis focused on the impact on cell proliferation and signaling effects of the HBx/PP2A PPI, it is likely that many HBV PPIs and network-identified HBV-associated genes also modify HCC behavior. Other HBV PPIs known to modify of tumor behavior that were significant after network analysis include the HBc targets AURKA and SRPK1, as well as the HBx targets HDAC1 and COPS4. As was the case for PP2A, while it is possible that HBV PPIs are targetable tumor drivers in HCC, these HBV PPIs may instead create targetable dependencies downstream of its PPIs. Future systems analysis may then uncover these dependencies, opening the door to new, mechanism-driven therapies for HCC.

## Methods

### Gene mutation analysis

To determine which protein coding genes were altered in the LIHC TCGA data set, we used the corresponding mutation data files and copy number calls on gene level provided by the Broad Institute TCGA GDAC. The mutation annotation file comprised 53,777 missense mutations as determined by Mutation Assessor (32) in 14,901 RNAs and 373 patients. We classified genes as altered (Mut) or wild type (WT) as follows: We removed variants classified as [‘Silent’, ‘IGR’, ‘5’UTR’, ‘3’UTR’, ‘5’Flank’, ‘3’Flank’, ‘RNA’, ‘Intron’], yielding 41,263 non-silent mutations across 13,675 genes. Additionally, we considered genes with amplifications or deletions as determined by GISTIC as mutated. Once mutations/CNAs were identified, we intersected them with the protein coding genes included in the ReactomeFI PPI reference network (https://reactome.org/).

As a result, we determined m=8,765 protein coding genes altered by either mutations, amplifications or deletions, in a set of n=366 patients. Patients with HBV and HCV co-infection were excluded. For the following analysis, we binarized this information for each gene: [0=WT, 1=Mut].

### Differential mutation analysis

To determine the effect of the viral infections on the mutational status of each gene in HCC, we assessed the differential mutation rates at gene level between:

A. HCV(+) and HCV(-) HCC cases.
B. HBV(+) and HBV(-) HCC cases.

For this purpose, we set up to map the inputs *x_ghepB_, x_ghepC_* to the output *y_g_*, where *g*∈{*g*_1_,…, *g_m_*}.The output *y_g_* is a one dimensional (1*d*) vector of length *n* representing the mutational status across the *n* HCC patients for each mutated gene *g* (0 = wild type; 1 = altered). The features *x_ghepB_* and *x_ghepC_* are 1*d* binary vectors of length *n* representing the viral infection status for HCV (1 = HCV(+); 0 = HCV(-)) -- *x_ghepC_*, and the viral infection status for HBV (1 = HBV(+); 0 = HBV(-)) -- *x_ghepC_*.

Given the response variable *y_g_* is binary, and we aimed to learn the impact of the viral infections on the mutation status, the first choice to formalize the problem was logistic regression. However, logistic regression returned perfect separation of the response which is a common problem in small sample size studies and imbalanced data. Perfect separation is accompanied by unstable regression coefficients, and can yield misleading findings. Since our aim was to estimate the risk of a mutation happening due to viral infection, and not to solve a binary classification problem, we used the solution proposed by Gelman et al. to obtain stable regression coefficients (33).

Following, we formally defined three Bayesian logistic regression models conditioned by independent Student-t prior distributions on the coefficients for each *g* out of the *m* mutated protein coding genes:

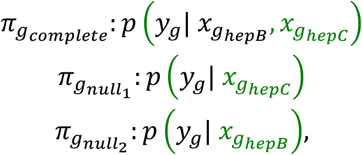

where *π_g_complete__* is defined by the probability mass function of the output *y_g_* given *x_ghepC_* and *x_ghepB_, π*_*g*_*null*_1___ is defined by the probability mass function of the output *y_g_* given *x_ghepC_*, and *π*_*g*_*null*_2___ is defined by the probability mass function of the output *y_g_* given *x_ghepB_* only. We used the default Cauchy distribution with mean 0 and prior scale 2.5 -- in the simplest scenario, a longer-tailed version of the distribution attained by assuming one-half additional success and one-half additional failure in a logistic regression.

To examine the impact of HBV infection on mutational status of HCC tumors, we compared the likelihood of the *π_g_complete__* model to the likelihood of each of the two alternative models. Hence, we calculated the deviances between the complete model and each of the two alternative models:

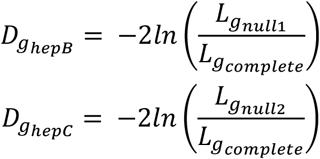

with *L* being the maximum likelihood, i.e. the probability of the data given the inputs *x_g_* and the parameter vector *θ* that maximizes 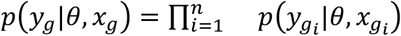, with *n* representing the number of samples.

### Affinity Purification – Mass Spectrometry

Affinity purification was performed as previously described (6,14). Briefly, HUH7 cells were transfected with FuGene 6 (Promega, Madison WI), and cell pellets were harvested 40 hours after transfection. Clarified cell lysates were incubated with prewashed Strep-Tactin beads (IBA Life Sciences) and allowed to bind for 2 hours. Following purification, complexes bound to beads were washed and then eluted with desthiobiotin (IBA Life Sciences). Proteins from cell lysates and AP eluates were evaluated by MS as below or separated by SDS-PAGE and either directly stained using the Pierce Silver Stain Kit (Thermo Fisher Scientific) or transferred to a PVDF membrane for immunoblotting.

AP eluates analyzed by MS underwent tryptic digest, desalting, and concentration and were then analyzed by LC/MS-MS on a Thermo Scientific Velos Pro Linear Ion Trap MS system or a Thermo Scientific Q Exactive Hybrid Quadrupole Orbitrap MS system equipped with a Proxeon EASY nLC II high-pressure liquid chromatography and autosampler system. Raw data were searched against SwissProt human protein sequences and the individual viral bait sequences using the Protein Prospector algorithm. PPIs were then scored with MiST algorithm using previously defined conditions (0.309 for reproducibility, 0.75 for specificity, and 0.006 for abundance). Cytoscape was used for visualization of the PPI network (34).

We also performed AP-MS on human bait proteins in the presence or absence of HBx. For these experiments, the bait proteins were cloned into the pcDNA4 vector with an N-terminal 3xFlag tag. These were transfected into HUH7 with pcDNA4-eGFP or pcDNA-ST-HBx, and then AP performed as above, but with anti-Flag-M2 magnetic beads (Sigma Aldrich, St. Louis MO). After washing, enriched proteins were eluted with Flag peptide and then subjected to MS as above. After acquisition, data were again searched against SwissProt human protein sequences. PPIs were scored with the MiST algorithm, as well as Saint (35) and Compass (36). A MiST score of 0.7, Saint BFDR <0.05 and Compass p value < 0.05 were used in combination to define a rigorous set of high-confidence PPIs; these were then overlaid onto the CORUM PPI map to define a strict PPI set. Label free protein quantities were determined for each prey using MaxQuant (37), and statistical inference perfromed with Ms Stats (38).

### Network propagation

We first separately propagated the HBV deviances -- *D_ghepB_*, through the reference network. We retained the propagated deviance scores in *S_d_g__*. Conceptually, *S_d_g__* indicates how likely it is that gene *g* is affected by proteins with differential viral-associated mutations. Thus, we estimated the viral effect on human proteins within the reference network by scoring the proximity of protein in the reference network to the HBV-interacting proteins at genomic level.

Next, we separately propagated the HBV MiST scores through the reference network. The propagated MiST scores were denoted by *S_p_g__* · *S_p_g__* represented the likelihood of gene *g* being affected by proteins that were physically interacting with viral proteins. Thus, we estimated the viral effect on proteins within the reference network, by scoring their proximity to the HBV-interacting proteins at physical level.

For network propagation, we used all human protein coding genes present in the reference network that were also expressed in LIHC (p = 9,803 proteins). Given the topology of the reference network, there are certain nodes (e.g. hubs) which will be ‘hot’ regardless of the initial scores represented by either deviances or MiST scores. To estimate the expected background of the propagated scores given the network topology, we performed 10,000 permutations in which we randomly reassigned the deviances *D_ghepB_*, and the HBV MiST scores. To calculate the significance of the propagation score of a specific gene, we ran the network propagation algorithm separately with the permuted deviances and MiST scores as input scores and we calculated empirical *p* values. The *p* values indicated how many times the propagated scores after permutation are greater than the real scores.

We used the gene-wise propagated MiST and deviances scores to calculate a combined measure of significance for each protein coding gene. We defined the combined significance score as:

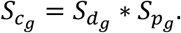

To obtain the null hypothesis distribution of the combined score given the network topology, we performed 10,000 permutations through which we randomly reassigned MiST scores and deviances. We applied the network propagation algorithm and calculated the product of the two propagated scores. We then calculated empirical *p* values corresponding to the combined score. The *p* values indicated which genes had network neighborhoods significantly enriched for both viral interactors and genes with differential mutation rates. We calculated the false discovery rate using the Benjamini–Hochberg procedure (48). The values represented the probabilities of erroneously finding genes with neighborhoods significantly enriched for both viral interactors and genes with differential mutation rates. Next, we used the interactions in the reference network between the proteins that showed combined significance and the viral-host interactions to build integrated interactomes of HBV in HCC (Figure 3C).

### Cell culture and treatments

Hep3B, HepG2, SNU-182, SNU-387, SNU-398, SNU-423, SNU-449, SNU-475, and PLC/PRF/5 were obtained from ATCC (Manassus, VA). HUH7 was obtained from the UCSF Cell Culture Facility and HUH6 was obtained from the RIKEN cell bank (Tsukuba, Japan). HLE, HLF, FOCUS and Hep40 were obtained from Dr. Ju-Seog Lee (MD Anderson Cancer Center). MHHC97-H and MHHC97-L were obtained from Zhongshan Hospital of Fudan University, Shanghai. Mycoplasma testing STR analysis were performed. Cells were cultured in DMEM with high glucose (Invitrogen, Carlsbad CA) with 10% fetal calf serum (Axenia BioLogIx, Dixon CA) for all experiments, with SNU-182, SNU-387, SNU-398, SNU-423, SNU-449, SNU-475 passaged in RPMI/10% FCS. Puromycin selections were performed at 1 μG/mL and hygromycin selections at 50 μG/mL. Cells were stably maintained in selection conditions.

Everolimus, LB-100, MG-132, MK-2206 and TAK-228 and were obtained from Selleckchem (Houston, TX), Cells were monitored periodically for mycoplasma contamination and identity confirmed with STR on receipt if not purchased from ATCC.

### Vectors and cloning

We used the following vectors to engineer cell models: pHR-SFFV-dCAS9-BFP-KRAB and pCRISPRia-v2 (BFP) and pCRISPRia-v2 (GFP) were kindly provided by Luke Gilbert (UCSF). Gateway compatible pLVX-TetOne-Puro was described previously (39), and adapted to hygromycin selection for use here. HBV genes were PCR subcloned from the 1.3wt HBV construct (40), obtained from Addgene, and all human proteins used in AP-MS were obtained from the Orfeome v8.1 (41).

### CRISPRi

Targeted CRISPRi analysis was performed following established protocols (42). Briefly, cells expressing dCas9-KRAB were sorted for BFP. dCas9-KRAB cells were then individually lentivirally infected with 4-5 sgRNAs against the gene target and selected with puromycin. Gene knockdown was confirmed by qPCR, and proliferation of CRISPRi cells was performed on a ZOOM Incucyte over 120 hours. Area under the curve (AUC) analysis of cell count at each time point was developed and compared to cells transduced with a non-targeting control (NTC) sgRNA.

### Mouse models

Wild-type (WT) FVB/N mice were obtained from Charles River Laboratories (Wilmington, MA) and *Rictor^fl/fl^* mice from The Jackson Laboratory (Sacramento, CA). Hydrodynamic injection was performed as previously (52). We injected 60μg pT3-EF1α-Cre or 60μg pT3-EF1α (empty vector control) together with 20μg pT3-EF1α-HA-myr-AKT and 20μg pT2-Caggs-RasV12 to delete *Rictor* while co-expressing AKT and Ras. Mice were housed, fed, and monitored in accord with protocols approved by the committee for animal research at the University of California, San Francisco (San Francisco, CA), and were monitored for signs of morbidity.

### Global phosphoproteomics

Huh7 cells with doxycycline-controlled 3xFLAG-HBx were treated with vehicle or doxycycline for 48 hours. Cells were then harvested in PBS, lysed in lysis buffer (8uM urea, 50mM Tris pH 8, 75mM NaCl, 1X protease and phosphatase inhibitors) and sonicated at 20% for 15 sec. Bicinchoninic acid (BCA) protein assay was performed to quantify protein lysates. Samples were reduced, alkylated and subjected to trypsin digestion at 37C overnight. Phosphopeptide enrichment was performed with immobilized metal affinity chromatography following established protocols (43). Enriched samples were analyzed on a Q Exactive Orbitrap Plus mass spectrometry system (Thermo Fisher Scientific). All mass spectrometry was performed at the Thermo Fisher Scientific Proteomics Facility for Disease Target Discovery at UCSF and the J. David Gladstone Institutes. Mass spectrometry data was assigned to human sequences and peptide identification and label-free quantification were performed with MaxQuant (version 1.5.5.1)(44). Data were searched against the UniProt human protein database (downloaded 2017). Statistical analysis was performed using the MSstats Bioconductor package (38). Phosphoproteomic data was uploaded to the PhosFate profiler tool (23) (Phosfate.com) to infer kinase activity.

### Multiplex inhibitor beads

Kinase chromatography, mass spectrometry and analytical processing were performed as described previously (24). Bait compounds were purchased or synthesized and coupled to sepharose. Cell lysates were diluted in binding buffer with 1 mol/L NaCl, and affinity purification was performed with gravity chromatography after preclearing. The bound kinases were stringently washed and then eluted with SDS followed by extraction/precipitation, tryptic digest and desalting. Liquid chromatography-tandem mass spectrometry (LC/MS-MS) was performed on a Velos Orbitrap (Thermo Scientific) with in-line high-performance liquid chromatography (HPLC) using an EASY-spray column (Thermo Scientific). Label-free quantification was performed with Skyline (45), and statistical analysis with Ms Stats (38).

### Analysis of public datasets

RPPA data was downloaded from UCSC Genome browser. Unpaired student’s t test was performed to assess the expression difference between the HBV and non-HBV group. Pearson correlation was used to compute the expression correlation between proteins.

### Western blots and antibodies

Cells were lysed with RIPA and Complete protease inhibitors and PhosStop phosphatase inhibitors following standard techniques. Blots were cut to allow analysis of multiple proteins at different molecular weights, and stripping was performed with Restore Western Blot stripping buffer (Thermo Fisher), to allow analysis of phosphorylation and total expression. All primary and secondary antibodies were obtained from Cell Signaling (Danvers, MA), except for actin and flag-HRP (Sigma, St. Louis, MO), HBx and SMC5 (Abcam).

### Quantitative Real Time PCR

RNA was extracted from cell lines with Trizol (Invitrogen) after washing with cold PBS to remove culture media. Reverse transcription was performed using the Superscript II RT kit (Invitrogen) with random hexamer primers (Roche) to produce cDNA from 2 micrograms total RNA. 18S and beta-actin were used as endogenous controls. All primers for analysis were generated with the Primer3 online tool (http://bioinfo.ut.ee/primer3); sequences available on request. Quantification was performed using SYBR green labeling.

### Immunohistochemistry

Analysis of mouse samples was performed on formalin fixed/paraffin embedded material with standard techniques and staining performed with anti-Ki67 (Thermo Fisher Scientific, #MA5-14520; 1:150), followed by DAB development with Vectastain Elite ABC.

---

**Figure S1:**
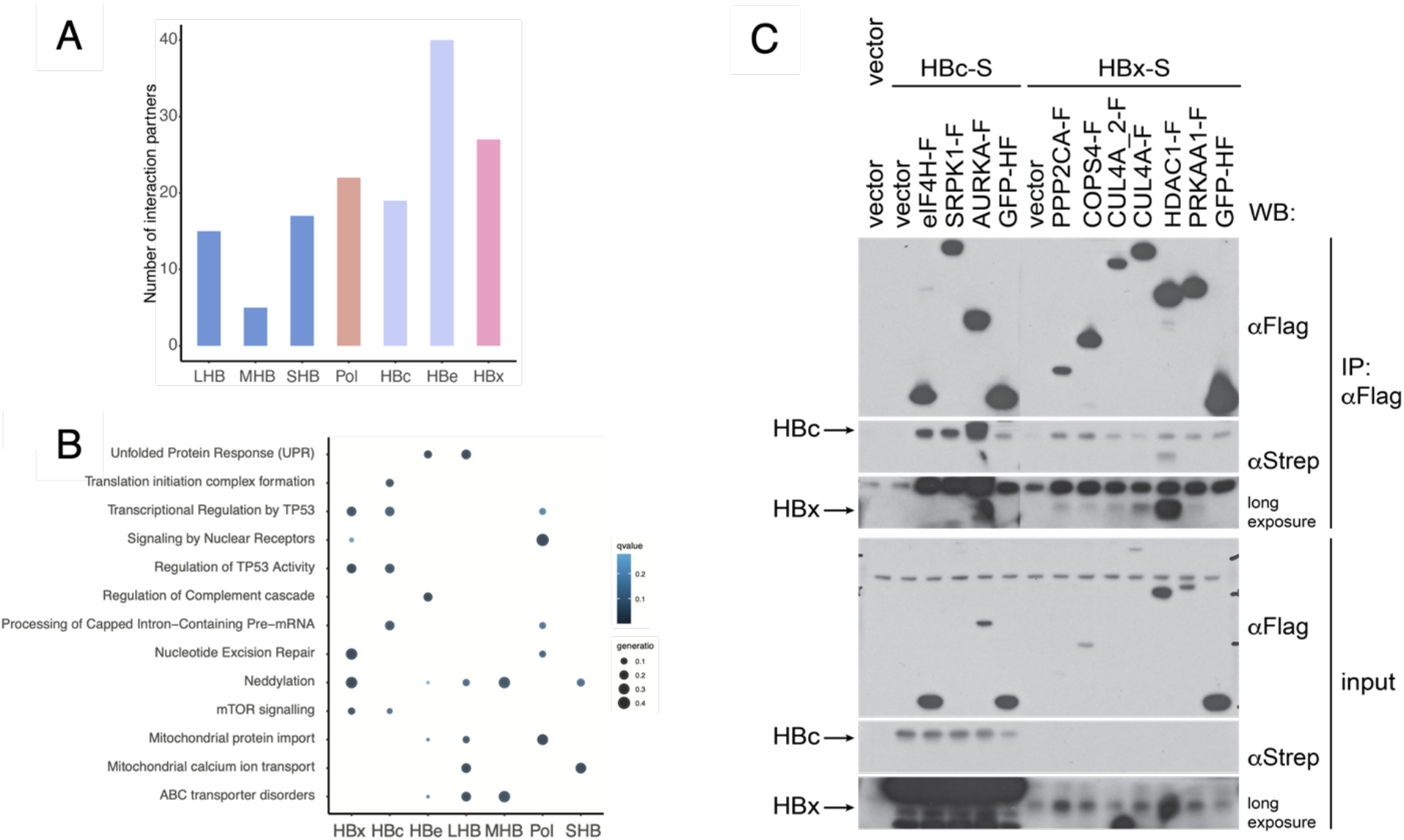
Direct confirmation of host/HBV PPIs: **A)** Number of high-confidence PPIs for each HBV protein. **B)** Selected results of bioinformatic analysis of HBV PPIs, assessing known HBV protein localization and functions, using databases such as KEGG and the NCI pathway interaction database. **C)** Selected HBc and HBx PPIs were cloned and flag-tagged, then co-expressed with either HBc-S or HBx-S to confirm AP-MS results using anti-FLAG IP. Empty vector and GFP are used as negative controls; n=3 biological replicates with 1 technical replicate per condition. Of note, HBc expression levels tended to be significantly higher than HBx expression levels, requiring longer exposures to visualize anti-STREP signal from HBx-S in either input or IP.

**Figure S2:**
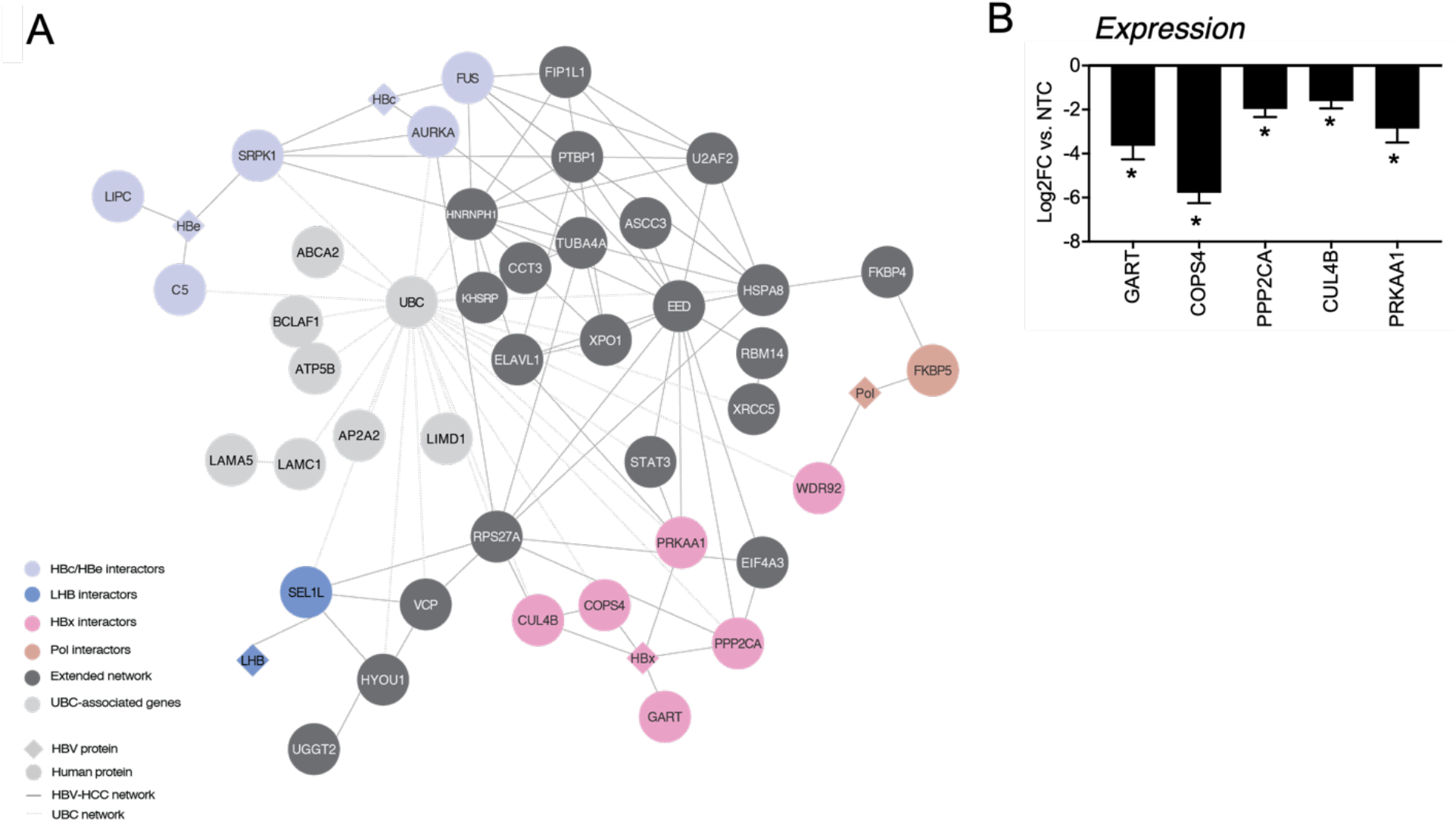
Network propagation to connect HBV PPIs and HCC mutations. **A)** Significant nodes from propagation analysis shown in network form, color-coded as in Figure 1C. An extended network of significant nodes from Reactome Fi PPIs that do not interact with an HBV protein is found, as well as a subnetwork of proteins that interact with UBC (ubiquitin), while significant based on HCC genetics, may be false positives for a contribution of protein interaction. **B)** Expression analysis of HBx PPIs that are significant from network propagation: HUH7 cells were engineered to express dCAS9-KRAB and then stably transduced with 4-5 sgRNA against each HBV PPI that reached significance in network analysis; the degree of knockdown is shown, p < 0.05 for all targets. n=1 biological replicates per sgRNA with a minimum of 3 technical replicates per sample.

**Figure S3:**
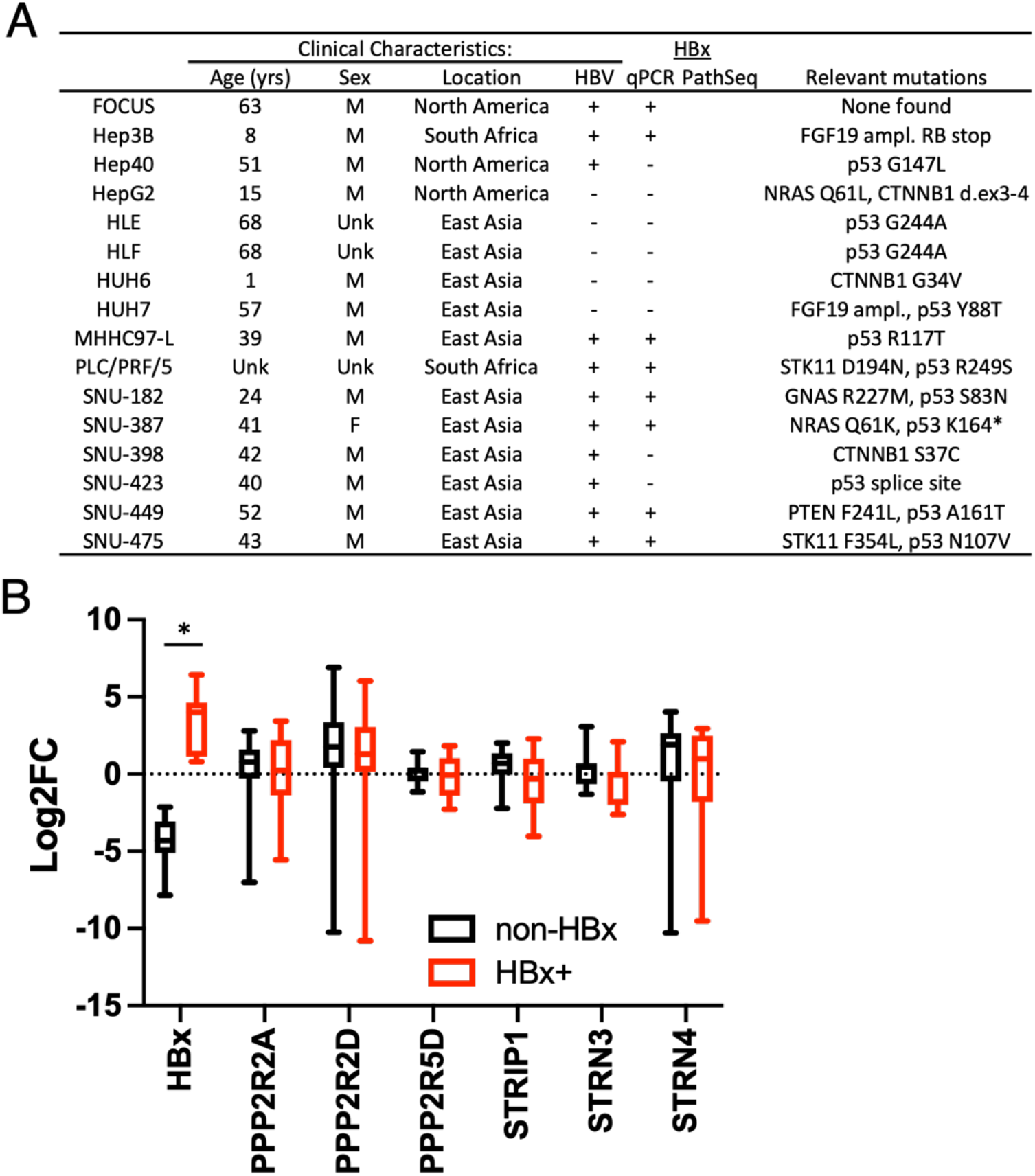
Additional PP2A analysis in HCC cell lines. **A)** Description of HCC cell panel: Clinical and genetic information describing cell lines analyzed, with HBx tested by qPCR. Clinical data extracted from the initial description of the lines, and genetic data from CCLE and other resources. **B)** Mean-centered expression of HBx and PP2A regulatory subunits across tumor cell line panel determined using QRT-PCR. No significant differences were seen except in HBx (* p<0.05); n=1 biological replicate with 3 technical replicates per condition. Of note, HBx classification corresponds to a PathSeq measurement of 0.4 HBV reads per million host reads.

**Figure S4:**
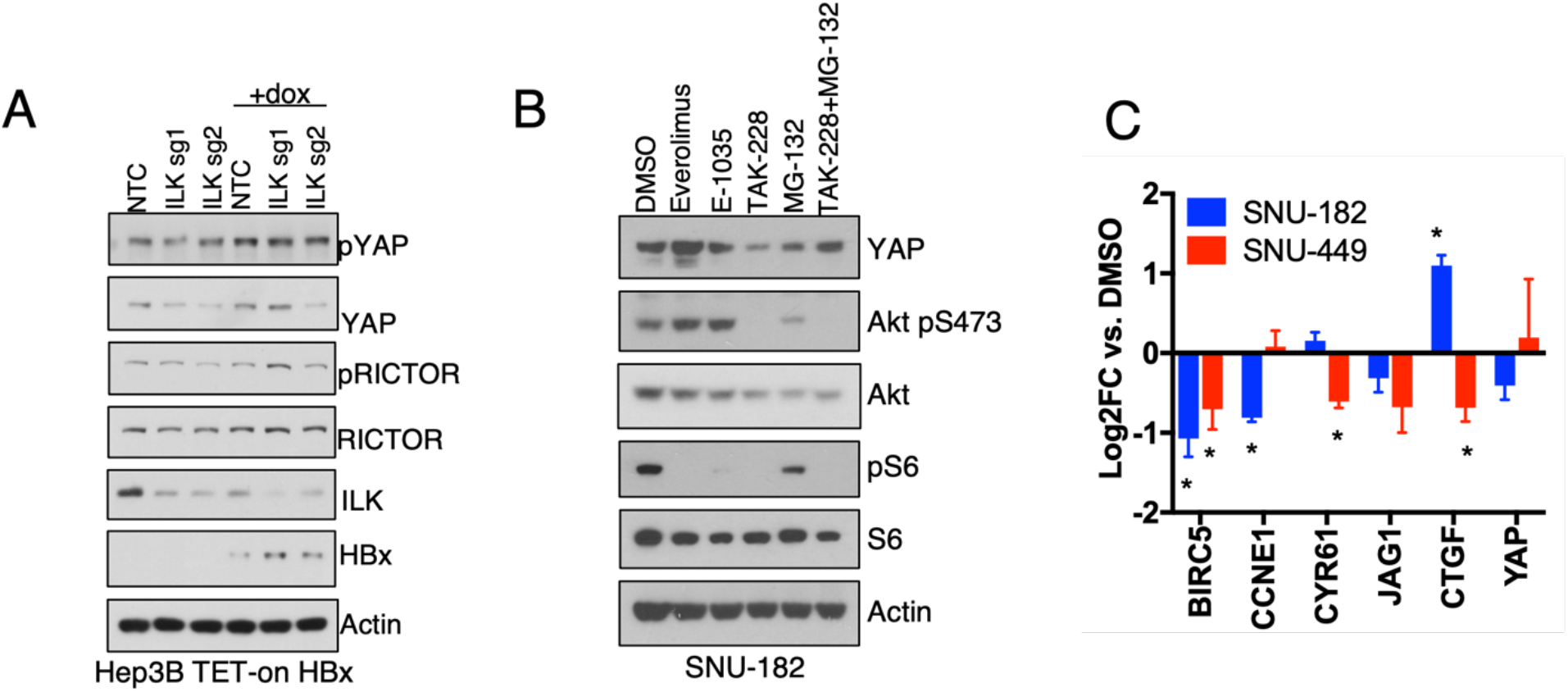
ILK and mTORC2 effects on YAP protein expression and transcriptional targets. **A)** Impact of ILK knockdown on YAP and RICTOR in Hep3B^dCAS9-KRAB^ tet-on HBx cells ± dox with non-targeting control (NTC) or two separate ILK-targeting sgRNA; n=3 biological replicates with 1 technical replicate per condition. **B)** Relative impact of mTORC1 and mTORC2 on YAP degradation: SNU-182 cells were treated for 4 hours with 100nM everolimus, 100nM of the Rapalink mTORC1 inhibitor E-1035, 100nM TAK-228, 10μM MG-132 or TAK-228+MG-132. n=3 biological replicates with 1 technical replicate per condition C) qPCR expression analysis of SNU-182 and SNU-449 cells of canonical YAP targets following 100nM TAK-228 treatment for 4 hours, normalized to DMSO treated cells; * p<0.05 as determined by t-test; n=3 biological replicates with 3 technical replicate per condition.

**Table S1:** HBV-HCC interactome

**Table S2:** Differentially mutated genes with HBV infection

**Table S3:** Integrated genomic interactome

**Table S4:** AP-MS of PP2A +/- HBx in HUH7 cells

**Table S5:** Global phosphoproteomics of HUH7 cells +/- HBx

**Table S6:** Phosfate analysis of differentially phosphorylated results from Table S5

**Table S7:** Multiplex Inhibitor Beads with mass spectrometry (MIB/MS) analysis in HUH7 cells

The data generated in this study are deposited in PRIDE will be made publicly available upon acceptance.

